# State-dependent modulation of activity in distinct layer 6 corticothalamic neurons in barrel cortex of awake mice

**DOI:** 10.1101/2021.11.08.467762

**Authors:** Suryadeep Dash, Dawn M. Autio, Shane R. Crandall

## Abstract

Layer 6 corticothalamic (L6 CT) neurons are in a strategic position to control sensory input to the neocortex, yet we understand very little about their functions. Apart from studying their anatomical, physiological and synaptic properties, most recent efforts have focused on the activity-dependent influences CT cells can exert on thalamic and cortical neurons through causal optogenetic manipulations. However, few studies have attempted to study them during behavior. To address this gap, we performed juxtacellular recordings from optogenetically identified CT neurons in whisker-related primary somatosensory cortex (wS1) of awake, head-fixed mice (either sex) free to rest quietly or self-initiate bouts of whisking and locomotion. We found a rich diversity of response profiles exhibited by CT cells. Their spiking patterns were either modulated by whisking-related behavior (∼28%) or not (∼72%). Whisking-responsive neurons exhibited either increases, activated-type, or decreases in firing rates, suppressed-type, that aligned with whisking onset better than locomotion. We also encountered responsive neurons with preceding modulations in firing rate before whisking onset. Overall, whisking better explained these changes in rates than overall changes in arousal. Whisking-unresponsive CT cells were generally quiet, with many having low spontaneous firing rates, sparse-type, and others being completely silent. Remarkably, the sparse firing CT population preferentially spiked at the state transition point when pupil diameter constricted and the mouse entered quiet wakefulness. Thus, our results demonstrate that L6 CT cells in wS1 show diverse spiking patterns, perhaps subserving distinct functional roles related to precisely timed responses during complex behaviors and transitions between discrete waking states.

**SIGNIFICANCE STATEMENT:** Layer 6 corticothalamic neurons provide a massive input to the sensory thalamus and local connectivity within cortex, but their role in thalamocortical processing remains unclear due to difficulty accessing and isolating their activity. Although several recent optogenetic studies reveal that the net influence of corticothalamic actions, suppression versus enhancement, depends critically on the rate these neurons fire, the factors that influence their spiking are poorly understood, particularly during wakefulness. Using the well-established *Ntsr1-Cre* line to target this elusive population in the whisker somatosensory cortex of awake mice, we found that corticothalamic neurons show diverse state-related responses and modulations in firing rate. These results suggest separate corticothalamic populations can differentially influence thalamocortical excitability during rapid state transitions in awake, behaving animals.

## INTRODUCTION

Layer 6 corticothalamic (L6 CT) neurons give rise to a massive descending projection that terminates in the thalamus. In sensory systems, these fibers outnumber ascending thalamocortical axons by 10:1, and CT axons contribute far more synapses than peripheral afferents (Sherman and Koch, 1986; Liu et al., 1995; Erisir et al., 1997; Deschenes et al., 1998; Van Horn et al., 2000; Winer et al., 2001). In addition, L6 CT cells send local axon collaterals to the more superficial layers of cortex that receive thalamic input (Gilbert and Wiesel, 1979; White and Keller, 1987; Zhang and Deschenes, 1997; Kim et al., 2014). Consequently, CT neurons can strategically influence thalamic throughput and cortical responsiveness.

Despite the anatomical prominence and a great deal of experimental effort focused on it (for review, see (Cudeiro and Sillito, 2006; Briggs and Usrey, 2008; Briggs, 2020)), a thorough understanding of CT functions remains elusive. This lack of clarity is partly due to the technical challenges of monitoring or selectively manipulating CT activity. However, recent advances in optogenetic tools and the *Ntsr1-Cre* mouse line (Gong et al., 2007), which expresses Cre recombinase in L6 CT cells, have helped overcome these limitations, accelerating inquiry into CT functions.

One key feature to emerge from these studies is that the net effect (suppressive or enhancing) generated by CT cells is likely to depend on the timing and rate at which they fire, with the sign and magnitude depending on their interactions with different excitatory and inhibitory circuit components (Olsen et al., 2012; Bortone et al., 2014; Mease et al., 2014; Crandall et al., 2015; Kim et al., 2015; Guo et al., 2017; Hasse and Briggs, 2017; Kirchgessner et al., 2020). However, a potential concern in using opsins to manipulate CT activity is that the uniform activation or suppression generated within the pathway may differ from naturally evoked patterns and thus affect thalamocortical activity differently. Furthermore, this approach ignores CT subtypes, which may have different *in vivo* activity profiles and functions (Tsumoto and Suda, 1980; Landry and Dykes, 1985; Sirota et al., 2005; Briggs and Usrey, 2009; Kwegyir-Afful and Simons, 2009). Thus, an open and unresolved question is whether CT neurons show variations in firing rates consistent with those necessary to modulate downstream targets and if their behavior varies across cells. This question is of particular interest, given most studies across modalities and species report that CT cells *in vivo* are silent and lack suprathreshold responses to sensory input in both anesthetized and awake animals (Tsumoto and Suda, 1980; Swadlow and Weyand, 1987; Swadlow and Hicks, 1996; Sirota et al., 2005; Kwegyir-Afful and Simons, 2009). In addition, because CT regulation is dynamic, state-dependent, and likely context-dependent (Steriade, 2001; Lee et al., 2008; Velez-Fort et al., 2018), it is critical to investigate these cells in awake, behaving subjects, something only a few studies have attempted (Beloozerova et al., 2003; Sirota et al., 2005; Augustinaite and Kuhn, 2020; Voigts et al., 2020; Clayton et al., 2021).

To determine the activity patterns of CT cells in behaving animals, we used single-cell juxtacellular recordings to study the spiking properties of optogenetically identified neurons in the whisker-related primary somatosensory cortex (wS1) of awake, head-fixed *Ntsr1-Cre* mice free to rest quietly or self-initiate whisking and locomotion. Rodents move their whiskers to gather information from their environment, and the activity patterns in wS1 contribute to tactile perception and sensorimotor processing (Diamond et al., 2008; Petersen, 2019). Yet no study has systematically recorded and functionally characterized the CT activity in wS1 during exploratory behavior. Consistent with recent calcium imaging and electrophysiological studies in visual and auditory cortices of awake behaving *Ntsr1-Cre* mice (Augustinaite and Kuhn, 2020; Clayton et al., 2021), we report that L6 CT neurons in wS1 are quiet, but a subset exhibit complementary activity patterns and spike rate modulation before and during active behavior and transitions between discrete waking states, suggesting general principles underlying CT activity across sensory modalities.

## MATERIALS AND METHODS

### Animals

All procedures were carried out in accordance with the National Institutes of Health (NIH) Guidelines for the Care and Use of Laboratory Animals and approved by the Michigan State University Institutional Animal Care and Use Committee (IACUC). The following mouse lines were used in this study: *Ntsr1-Cre* (MMRRC: 017266-UCD) (Gong et al., 2007), Ai32 (Jackson Labs: 024109) (Madisen et al., 2012), Ai14 (Jackson Labs: 007908) (Madisen et al., 2010), and Crl:CD1 (ICR: Charles River: 022). All *Ntsr1-Cre* mice used had Crl:CD1 genetic backgrounds. For most experiments, homozygous *Ntsr1-Cre* mice were crossed with a floxed ChR2 reporter line (homozygous Ai32), resulting in heterozygous experimental mice for the indicated genes. For viral injections, homozygous *Ntsr1-Cre* mice were crossed with Crl:CD1 mice. Experimental mice were group-housed with same-sex littermates before surgery and singly housed post-surgery in a dedicated animal care facility maintained on a reversed 12:12 h light-dark cycle. Food and water were available *ad libitum*. Both male and female mice were used in this study.

### Surgical Procedures and Training

Mice were 6-60 weeks old at the time of surgery. Briefly, mice were anesthetized with a Ketamine-Dexdomitor mixture diluted in sterile saline (KetaVed, 100 mg/kg; Dexdomitor, 0.25 mg/kg; intraperitoneally). Once deeply anesthetized, the eyes were covered with a thin layer of ophthalmic ointment to prevent drying (Patterson Veterinary Artificial Tears). For chronic electromyography (EMG) recordings, fine Teflon-coated tungsten microwires (50 μm: A-M Systems # 795500) were first inserted under the skin into the left mystacial pad musculature, as previously described (Fee et al., 1997; Berg and Kleinfeld, 2003). The distal tips of wires (∼1 mm) were stripped of insulation and hooked for stable positioning within the pad. The EMG wire was then positioned on the head before the incised skin was closed with a surgical adhesive (GLUture). Mice were then placed into a digital stereotaxic frame with an integrated warming base that maintained core body temperature (Stoelting). Next, the skull was exposed using fine surgical scissors, the periosteum overlying the skull removed, and a small craniotomy (∼0.75 mm diameter) made over the right wS1 (coordinates relative to bregma: 3.4 mm lateral and 1.1 mm posterior). A custom-built titanium head-post was then affixed to the skull with adhesive luting cement (C&B Metabond), and the resected skin closed around the outside of the head-post with tissue adhesive (GLUture). The EMG wire was attached to a gold pin (A-M Systems) with cold solder and secured in the luting cement. The craniotomy was then covered with Kwik-Cast (WPI), and Buprenorphine (0.05-0.1 mg/kg) was injected subcutaneously for postoperative pain relief. Mice were also given Antisedan (2.5 mg/kg) to reverse the effects of Dexdomitor. After surgery, mice were allowed to recover on a heating pad for 1-2 hours before returning to their home cage.

After 48-72 hours of recovery, implanted mice were handled and habituated to head fixation within the recording setup. We clamped the head-post to metal posts, positioned the mouse on a flexible circular treadmill, and then allowed mice to stand/walk/run freely in this position. Mice acclimated quickly to head fixation over the course of two 30-min sessions and ran naturally on the wheel, never showing signs of distress. Physiological recordings began once the animal was accustomed to head fixation, which was ∼72 hours after surgery. All behavioral training and physiological recording were performed during the dark phase.

### Stereotactic Injections

For some mice (n = 2), an adeno-associated virus (AAV) that encoded Cre-dependent genes for hChR2(H134R)-EYFP fusion proteins was injected into the right wS1 during the head-posting procedure, as previously described (Crandall et al., 2015; Martinetti et al., 2022) (pAAV1.Ef1a.DIO.hChRr2(H134R)-eYFP.WPRE.hGH, Addgene: 20298-AAV1). Briefly, after the craniotomy was made over wS1, a small volume of the virus was then pressure-ejected via a glass micropipette attached to a Picospritzer pressure system (0.33 μl over 10-15 min; titer = ∼3.4 x 10^12^ viral genomes/ml). Viral solutions were injected between 0.8-1.0 mm ventral to pia surface. Following injection, the pipette was held in place for 10 min before withdrawal from the brain. After the procedure, mice were allowed to recover and express the viral vector for 10-12 days.

### In Vivo Awake Recordings and Data Acquisition

On the day of recording, Kwik-cast was gently removed from the skull, exposing the craniotomy, and a drop of saline (0.9% NaCl) was placed on the skull to keep the exposed craniotomy moist. *In-vivo* juxtacellular recordings were achieved using borosilicate glass pipettes with tip resistances of 3-6 MΩ (tip diameter: 2-3 µm) when filled with HEPES-buffered artificial cerebrospinal fluid (ACSF) containing (in mM): 126 NaCl, 3 KCl, 5 HEPES, 1.25 NaH_2_PO_4_, 1 MgSO_4_, 1.2 CaCl_2_, 26 NaHCO_3_, and 10 glucose, pH 7.25 with NaOH, osmolarity: 300 mOsm). Recording pipettes were mounted on a four-axis micromanipulator (Sutter Quad) and lowered through the craniotomy at a normal angle, avoiding blood vessels with guidance by stereomicroscope (Leica MZ10F). At a depth of 200-400 μm, the penetration was temporarily paused, and the craniotomy and pipette stabilized by applying warmed Agar or Agarose (3%) in saline (0.9% NaCl). The initial speed of advancement was 5-10 µm/s, but manipulator advancement was slowed down to 2-4 µm/s at a depth of 800 µm until a cell was encountered. During the search for a single cortical neuron, physiological activity and electrode resistance were monitored using Cambridge Electronic Design (CED) hardware and software and an audio amplifier (A-M Systems Model 3300). Upon detecting spontaneous or light-evoked (see below) small extracellular spikes, the recording pipette was slowly advanced until we could achieve a juxtacellular recording configuration, defined as a recording with spikes having a positive polarity initial phase and an increase in electrode resistance just before, or coincident with, the detection of a neuron. Juxtacellular recordings were considered for analysis if action potential amplitudes were ≥ 1 mV, remained stable throughout the recording, and if the series resistance varied ≤ 20%. The final data set also included six neurons with extracellular spiking. The exclusion of these neurons from the dataset had no qualitative impact on the interpretation of the results.

All electrophysiological and behavioral signals were acquired and digitized at 20 kHz using Cambridge Electronic Design hardware and software (Power 1401-3 and Signal 6). Juxtacellular data was obtained using a Multiclamp 700B amplifier (Molecular Devices), and signals were low pass filtered at 10 kHz before digitizing. During juxtacellular recordings, the pipette capacitances were neutralized, and series resistances (typically 10-25 MΩ) were compensated online. Series resistances were continually monitored during experiments to ensure sufficient compensation. The EMG electrode was connected to an A-M Systems preamplifier head-stage (20x) through a connector with small input leads, and then the voltage signals were further amplified 100x and band-pass filtered between 10 Hz to 3kHz using an A-M Systems 3600 amplifier. Pupil signals were obtained from the eye contralateral to the recording site using a dedicated infrared (880 nm) eye-tracking system at a sampling rate of 120 Hz (ISCAN ETL-200 rodent series), and the output was interfaced via Cambridge Electronic Design hardware (Power 1401-3). An additional white LED was positioned above the mouse and provided low-intensity illumination such that the pupil was approximately mid-range in diameter when the animal was calm but alert. All recordings were performed in isoluminant conditions without any change in the ambient light during a recording session. Locomotion epochs were tracked using an infrared emitting LED (940 nm) and an infrared-sensitive phototransistor (Sparkfun QRE1113). Locomotion was detected when emitted light was interrupted or reflected by a black and white pattern printed on the side of the wheel.

### In vivo Photostimulation and Identifying ChR2-expressing Neurons

We used local photostimulation to identify ChR2-expressing L6 CT neurons for *in vivo* recordings during awake behavior (Lima et al., 2009; Moore and Wehr, 2013; Munoz et al., 2014; Munoz et al., 2017; Yu et al., 2019). To couple light stimulation with juxtacellular recordings, we used a two-port glass pipette holder to insert an optical fiber into the recording pipette, providing direct illumination through the tip (A-M Systems, Optopatcher) (Katz et al., 2013). The internal optic fiber (0.22 NA, inner diameter of 200 μm, 1.25 mm OD ceramic ferrule, Thorlabs, FG200UCC) was positioned as deep as possible in the pipette to achieve maximum light intensity at the tip of the pipette (3-8 mW). The internal optic fiber was coupled to a 465 nm LED light source (Plexon, 94002-002) by a patch cable (0.22 NA, inner diameter of 200 μm, 1.25 mm outer diameter ceramic ferrule, Thorlabs, M86L01) and driven by a single channel LED driver (Plexon, 51382). LED on/off times were fast (< 50 μs) and of constant amplitude and duration when verified with a fast photodiode (Thorlabs, DET36A).

While advancing the recording pipette, we used brief LED pulses (5 ms, Max LED intensity) delivered at a low frequency (2 Hz) to identify putative ChR2-expressing neurons. Once a neuron was encountered and stable under the recording pipette, we ran a block of photostimulation trials (5 ms pulse duration, max intensity, >50 repetitions) to establish if the neuron expressed ChR2. After the photostimulation block, we recorded the neural activity simultaneously with other time-varying signals like EMG, pupil diameter, and locomotion. To confirm ChR2-expressing CT neurons offline, we focused our analysis on two characteristics: (1) the probability of light-evoked spikes and (2) the latency of peak light-evoked spikes.

### In Vivo Juxtacellular Labeling

In a subset of experiments, we filled the juxtacellular recording pipette with HEPES-buffered ACSF containing 2.5% Neurobiotin (Vector Laboratories) so that the recorded cell could be labeled and subsequently identified (Pinault, 1996). Following the recording, we injected 5-15 nA positive current pulse (200-400 ms, 0.25 Hz) through the recording pipette to modulate the spiking activity. We continually monitored evoked spikes throughout the labeling protocol, and current amplitudes were adjusted to avoid increased spontaneous activity and cell damage. The recording pipette was slowly retracted once the labeling protocol was complete (typically 300-500 current pulses with evoked spikes). Following a survival period of 2-24 hrs, mice were sacrificed and brain tissue fixed for subsequent histology.

### Histology and Imaging

All tissue for histology was prepared from acute brain slices, as previously described (Crandall et al., 2017; Martinetti et al., 2022). Briefly, animals were anesthetized with isoflurane before being decapitated. Brains were removed and placed in a cold (∼4°C) oxygenated (95% O_2_, 5% CO_2_) slicing solution containing (in mM) 3 KCl, 1.25 NaH_2_PO_4_, 10 MgSO_4_, 0.5 CaCl_2_, 26 NaHCO_3_, 10 glucose, and 234 sucrose. Brain slices (300-1000 µm thick) containing wS1 were cut using a vibrating tissue slicer (Leica VT1200S) and immediately transferred to a 4% paraformaldehyde in 0.1 M phosphate buffer solution overnight at 4°C (18 – 24 hr). The next day, slices were changed to a 30% sucrose in 0.1 M phosphate buffer solution until re-sectioned (4°C; 2 – 3 days). Individual brain slices were re-sectioned at 100 µm using a freezing microtome (Leica SM2010 R). Floating sections were washed twice in 0.1 M phosphate buffer followed by 3 washes in 0.1 M phosphate buffer with 0.15 M NaCl, pH 7.4 (PBS, 5 min per wash). After washing, sections were incubated for 1 hour at room temperature in a blocking solution containing 0.1% Tween, 0.25% Triton X-100, 10% normal goat serum in PBS. Sections were then incubated using Streptavidin/Biotin Blocking Kit (Vector Labs, Cat # SP-2002), 30 minutes in streptavidin solution, followed by 30 minutes in biotin solution with a brief rinse of PBS after each. Sections were then incubated with Alexa Fluor 568-conjugated streptavidin (Thermo-Fisher Scientific, Cat #S11226, 1:5000, 2μg/μl) solution made up in blocking solution for 3 hours with rotation at room temperature. Following incubation, sections were washed 3 times in PBS, then 2 times in 0.1 M phosphate buffer solution (10 min per wash), mounted, and cover-slipped using Vectashield with DAPI (Vector Laboratories H-1500). Fluorescent conjugates of streptavidin were stored and prepared as recommended by the manufacturer. Confocal image stacks of labeled neurons were taken using an Olympus FluoView 1000 filter-based Laser Scanning Confocal Microscope with a UPlanSApo (NA 1.40) 100x oil immersion objective and updated Version 4.2 software (laser excitation 405nm, 514nm, and 543nm). Image brightness and contrast were adjusted offline using Fiji software (Schindelin et al., 2012). Three blinded independent observers examined cells to determine the presence or absence of membrane-bound ChR2-eYFP associated with the *in vivo* filled neurobiotin cells.

### Electrophysiological Data Analysis

Electrophysiological and behavioral data were analyzed in Spike 2 (CED) and Matlab (Mathworks; Natick, MA). After the experiment, juxtacellular spiking data was high pass filtered (250-270 Hz, finite impulse filters (FIR) in Spike 2), and single action potentials (APs) were detected offline using Spike 2 software (CED). The spike identification procedure involved a two-step process: first setting a threshold with two trigger points (removing noise from the putative spike) and then selecting a template from the recorded spike waveforms. Spike shapes with 60% of the points matching the pre-selected template were captured and added, and their event times saved. We then converted the timestamps of identified spikes and all other electrophysiological and behavioral data to Matlab files, and all subsequent analyses were performed using custom-written scripts in Matlab.

Briefly, spike timestamps were used to calculate the continuous instantaneous firing rate of the spike train. The instantaneous firing rate was calculated as the inverse of the inter-spike intervals and then smoothed with a low pass Butterworth filter of 100 ms window size, which was applied in both the forward and backward directions to ensure no net smearing of data in a particular direction. Next, the mystacial pad EMG signal was rectified and filtered with a 2^nd^ order Butterworth filter with a desired final cut-off frequency of 5 Hz and applied in both forward and backward directions to avoid temporal smearing. The final signal was a continuous trace of the rectified EMG activity envelope during a recording session. The raw pupil diameter signal was also smoothed with a low pass Butterworth filter with 100 ms window size, running forward and backward. Finally, the signal of the infrared phototransistor was rectified, differentiated, and low-pass filtered (5 Hz) to get a smoothed locomotion signal.

The onset of locomotion and EMG epochs were calculated based on a threshold above the preceding baseline signal. All locomotion epochs, whisking epochs with or without locomotion, were initially detected manually. Once identified, we calculated the mean and standard deviation of the baseline as the average of the preceding 1 s of respective signals. We then defined locomotion onset objectively as the point +/- 500 ms around the manually detected onset that exceeded the baseline plus five times the standard deviation. Similarly, we defined whisking/EMG onset as the point exceeding baseline plus three times the standard deviation.

We extracted 5 s of data before and after each detected whisking/EMG onset for subsequent analysis of whisking with concomitant locomotion (whisking/locomotion) epochs. For a recorded neuron to be included for analysis, at least three locomotion epochs were needed. For whisking only (whisking) epochs, we extracted 3 s before and after each detected whisk. Every neuron considered in this study had greater than 10 such events. For statistical analysis of within neuron whisking-related modulation, the baseline period was defined as -4 s to -1 s before whisking onset for whisking/locomotion epochs and -1.5 s to -0.5 s before whisking onset for solitary whisking episodes. The whisking period was defined as 3 s and 2 s starting from whisking onset for whisking/locomotion epochs and for solitary whisking epochs, respectively. For a neuron to be considered whisking-responsive, significant modulation (p < 0.05, Wilcoxon signed-rank test) from baseline for either whisking with or without locomotion epochs was needed.

To determine whether whisking-responsive CT neurons were better correlated with whisking or a change in arousal, we cross-correlated instantaneous spiking activity with the time course of either the EMG signal or pupil dynamics. For whisking/locomotion epochs, we used 4 s of data centered on the whisking onset, whereas for solitary whisking, we used 3 s of data centered on whisking onset for cross-correlations. The analysis yielded correlation coefficients (r-value) at every time bin (of 1 ms) for lag and lead of 1 s each.

### Statistical Analysis

Statistical comparisons were performed in Matlab. Details of specific comparisons are provided in the results section. Briefly, non-parametric tests were held as appropriate based on a test of normality using the Shapiro-Wilk test. Paired and unpaired comparisons were performed using the Wilcoxon signed-rank test and the Mann-Whitney tests, respectively. All time-varying data traces (average instantaneous spike rates, EMG profile, and pupil dynamics) are represented as mean ± SEM, and any numerical representation (e.g., average spiking during a fixed time window and mean r-value) is represented as mean ± SD, and statistical significance was defined as p < 0.05. However, exact p-values are provided wherever applicable.

## RESULTS

### Optogenetic tagging and functional identification of L6 CT neurons in wS1

To investigate the behavior of L6 CT cells in awake, behaving mice, we used an *in vivo* Channelrhodopsin (ChR2)-assisted patching strategy to target individual genetically identified CT neurons for juxtacellular recording (Lima et al., 2009; Moore and Wehr, 2013; Munoz et al., 2014; Munoz et al., 2017; Yu et al., 2019). For these experiments, we took advantage of the *Ntsr1-Cre* driver mouse line (Gong et al., 2007), which selectively labels L6 CT neurons in sensory cortices, including wS1 (Bortone et al., 2014; Kim et al., 2014; Crandall et al., 2015; Guo et al., 2017). Consistent with previous reports, we found that *Ntsr1-Cre* positive somata in wS1 were restricted to L6 and that their axons/terminals were present in whisker-related nuclei of the somatosensory thalamus (**Fig. 1*A***). To render genetically defined CT neurons responsive to light, we conditionally expressed the light-sensitive cation channel ChR2 by either crossing the *Ntsr1-Cre* driver mouse to a reporter mouse line (Ai32) that conditionally expresses ChR2-enhanced yellow fluorescent protein (ChR2-EYFP) or by viral transduction with a Cre-dependent adeno-associated virus expression vector (AAV) (Madisen et al., 2012; Crandall et al., 2015). To directly relate CT activity to behavior and waking state, we recorded individual cells located across the depth of L6 in awake, head-fixed mice positioned on a flexible wheel-apparatus while simultaneously monitoring the mystacial-pad EMG for whisking, pupil diameter for changes in general arousal, and wheel movement for locomotor bouts (**Fig. 1*B*, left**). Previous studies have shown pupil diameter to be a reliable index of arousal state in mice (Reimer et al., 2014; McGinley et al., 2015a; Vinck et al., 2015). For identification of L6 CT cells *in vivo*, we used brief focal LED stimuli (465 nm, 5 ms, 2 Hz, 3-8 mW) directed through the glass recording pipette to stimulate ChR2-expressing CT cells effectively at the tip of the pipette (**Fig. 1*B*, right**).

**Figure 1.**
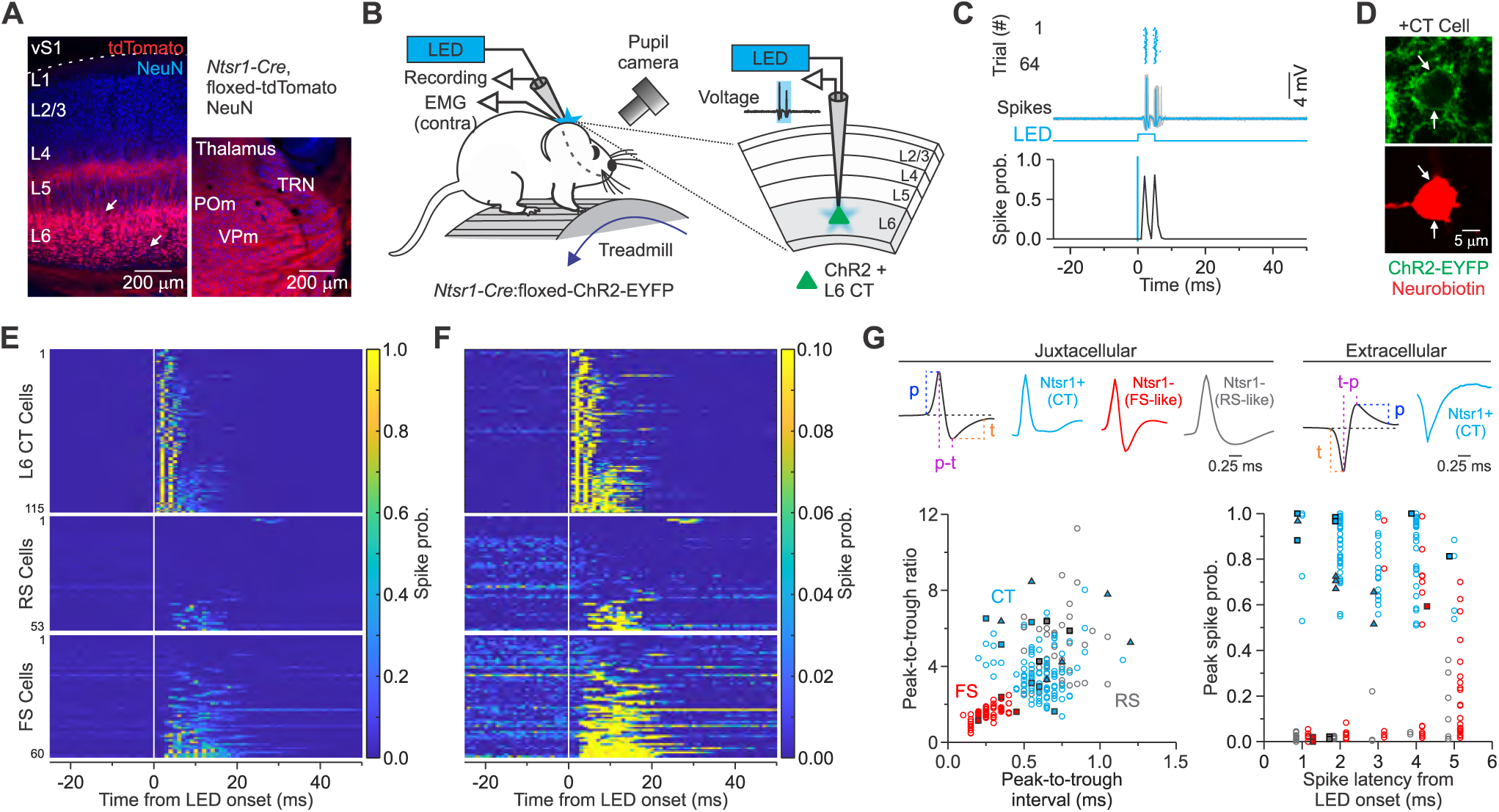
Optogenetic tagging and functional identification of genetically defined L6 CT neurons in wS1 of the *Ntsr1-Cre* driver mouse line. ***A,*** Left: Confocal image of a fixed brain section (40 μm), centered on wS1, from an *Ntsr1-Cre* x floxed-tdTomato reporter (Ai14) mouse. White arrows indicate Ntsr1 expressing CT somata in L6. Right: Confocal image from the same fixed section, centered on the somatosensory thalamus, showing tdTomato-labeled CT axons terminating densely (VPm: ventral posterior medial nucleus; POm: posterior medial nucleus; TRN: thalamic reticular nucleus). The tissue was stained immunohistochemically for NeuN (blue). ***B,*** Schematic of the experimental setup. Head-fixed *Ntsr1-Cre* x floxed-ChR2-EYFP reporter (Ai32) mice stood or walked/ran on a flexible circular treadmill during juxtacellular recordings from neurons in wS1 while simultaneously monitoring mystacial pad EMG, pupil diameter, and locomotion. We used an optical fiber located inside the glass recording pipette to deliver blue light locally through the tip for stimulation and identification of ChR2-expressing L6 CT cells *in vivo* (optogenetic tagging; 465 nm LED). ***C,*** Top: Raster plot of light-evoked spiking (blue dots) order by trial number for a ChR2-expressing L6 CT neuron. Brief LED flashes (5 ms) from the pipette tip evoked reliable spikes with short onset latencies (blue voltage trace) across all trials (grey traces). Bottom: Peristimulus time histogram (PSTH) of spike probability for 64 trials aligned on LED onset (blue line) (bin size: 1 ms). We observed reliable light-evoked spikes within 2 ms of LED onset for this neuron. ***D,*** Post hoc confocal images of the same neuron (shown in C) labeled with neurobiotin by *in vivo* juxtacellular methods during the recording. Membrane ChR2-EYFP expression histologically confirmed that this recorded cell was a ChR2-expressing L6 CT neuron in the *Ntsr1* x Ai32 mouse. ***E,*** *Top:* The normalized spike probability of all optogenetically tagged L6 CT neurons during brief photostimulation (n = 115 cells from 27 mice), sorted by mean spike probability in the 5 ms after LED onset. The white line indicates LED onset. *Middle and Bottom:* The normalized spike probability of all RS (n = 53; peak-trough duration *≥* 0.5ms) and FS cells (n = 60; peak-trough duration ≤ 0.4ms, peak-trough ratio ≤ 2.5), respectively. ***F,*** Same as E but with a different scale (0-0.1 normalized spike probability) for better visualization of spike latency and suppressed units. ***G,*** *Top Left:* Waveforms of spikes (recorded juxtacellularly) of a photoactivated L6 CT neurons (blue), putative fast-spiking (FS) interneuron (red), and regular-spiking (RS) cell (grey). Top Right: Example waveform of a spike recorded extracellularly (n = 6 of 115 cells) from a photoactivated L6 CT cell (blue). Insets show parameters measured. Bottom left: Plot showing the separation of putative FS interneurons (red) and RS cells (grey) from optically identified L6 CT (blue) based on the peak(p)-to-trough(t) duration and the peak-to-trough ratio of their average recorded spike waveform. Right: Plot showing the magnitude and timing of peak spike probability during photostimulation. Light-evoked response of L6 CT cells exhibited greater spike probability and shorter latency to peak spiking probability than putative FS interneurons and RS cells. The squares indicate cells confirmed based on neurobiotin staining and EYFP expression (6 CT cells and 6 non-CT cells), whereas the triangles represent the 6 extracellular CT units included in the dataset. The trough-to-peak ratio and interval were plotted for all extracellularly recorded CT cells. The absolute peak-to-trough ratio was plotted for all recorded cells.

**Figure 1C** shows an example of a photo-activated neuron identified as a CT cell based on the characteristics of its light-evoked activity. This neuron had a cumulative spike probability of one within the first 3 ms of LED onset. Following the recording, we labeled the cell with neurobiotin by juxtacellular iontophoresis and confirmed ChR2 expression histologically (Pinault, 1996) (**Fig. 1*D***). We observed similar light-evoked activity in 6 cells confirmed posthoc expressing ChR2 (**Fig. 1*G***). Across the entire population, identified L6 CT cells had a spike probability of 0.5 or greater within 3 ms of LED onset and a cumulative probability of 1.0 or greater within 5 ms of LED onset (defined as the sum of the spike probability over the first five 1-ms bins) (n = 115 cells from 27 mice; 17 males and 10 females) (**Fig. 1*E-G***). We performed additional analyses on spike waveform components of all light-insensitive cells to separate putative FS neurons and regular spiking (RS) neurons from our CT population. The peak spike probability within 5 ms of LED onset of all RS (Mean ± Std. Dev. = 0.03 ± 0.07; n = 53 cells; peak-trough duration ≥ 0.5 ms) and most FS cells (Mean ± Std. Dev. = 0.85 ± 0.15; n = 60; peak-trough duration ≤ 0.4ms, peak-trough ratio ≤ 2.5) was lower than that of CT cells (**Fig. 1*E-G***). Putative FS and RS cells also had a longer latency to peak spike probability from LED onset than CT neurons (Mean ± Std. Dev. = 2.85 ± 1.61 ms; **Fig. 1*E-G***). Consistent with these observations, we found that histologically recovered cells with a low spike probability (< 0.5) or a significant delay to light-evoked responses (> 5 ms) were negative for ChR2 expression (n = 6 cells; **Fig. 1*G***). Thus, high spike probability and short-latency to peak spike probability are reliable physiological criteria for identifying ChR2-expressing L6 CT neurons *in vivo*.

### Four broad classes of L6 CT cells based on their activity patterns in awake mice

We recorded the activity of all L6 CT neurons encountered by our pipette (n = 115) during simple free behavior, in which the head-fixed mouse was allowed to rest quietly or self-initiate whisking-related behavior. Based on the measured EMG signal and recorded wheel movement, we defined the behavioral state of the mice as 1) quiet wakefulness with no whisker movement, 2) whisking, and 3) whisking with concomitant locomotion (whisking/locomotion).

The activity patterns differed considerably among CT neurons, even within a single electrode penetration (**Fig. 2**). We found that most L6 CT cells did not show significant changes in spike rate during episodes of whisking or whisking/locomotion (Whisking-unresponsive: 72%; 83 of 115 CT neurons; mean spiking rate change around whisking onset; p > 0.05; Wilcoxon signed rank test; **Fig. 2*A,C,D***). Of these Whisking-unresponsive CT cells, some were spontaneously silent, firing no spikes during the entire recording session (Silent-type: 39.8%; 33 of 83 Whisking-unresponsive neurons), whereas the majority had low spontaneous firing rates, typically less than 0.5 spikes/s (Sparse-type: 60.2%; 50 of 83 Whisking-unresponsive neurons; mean spontaneous rate: 0.22 ± 0.55 spikes/s). These data are consistent with results from other species and sensory cortices showing many L6 CT neurons are silent or sparsely firing (Tsumoto and Suda, 1980; Swadlow, 1994; Beloozerova et al., 2003; Sirota et al., 2005; Stoelzel et al., 2017).

**Figure 2.**
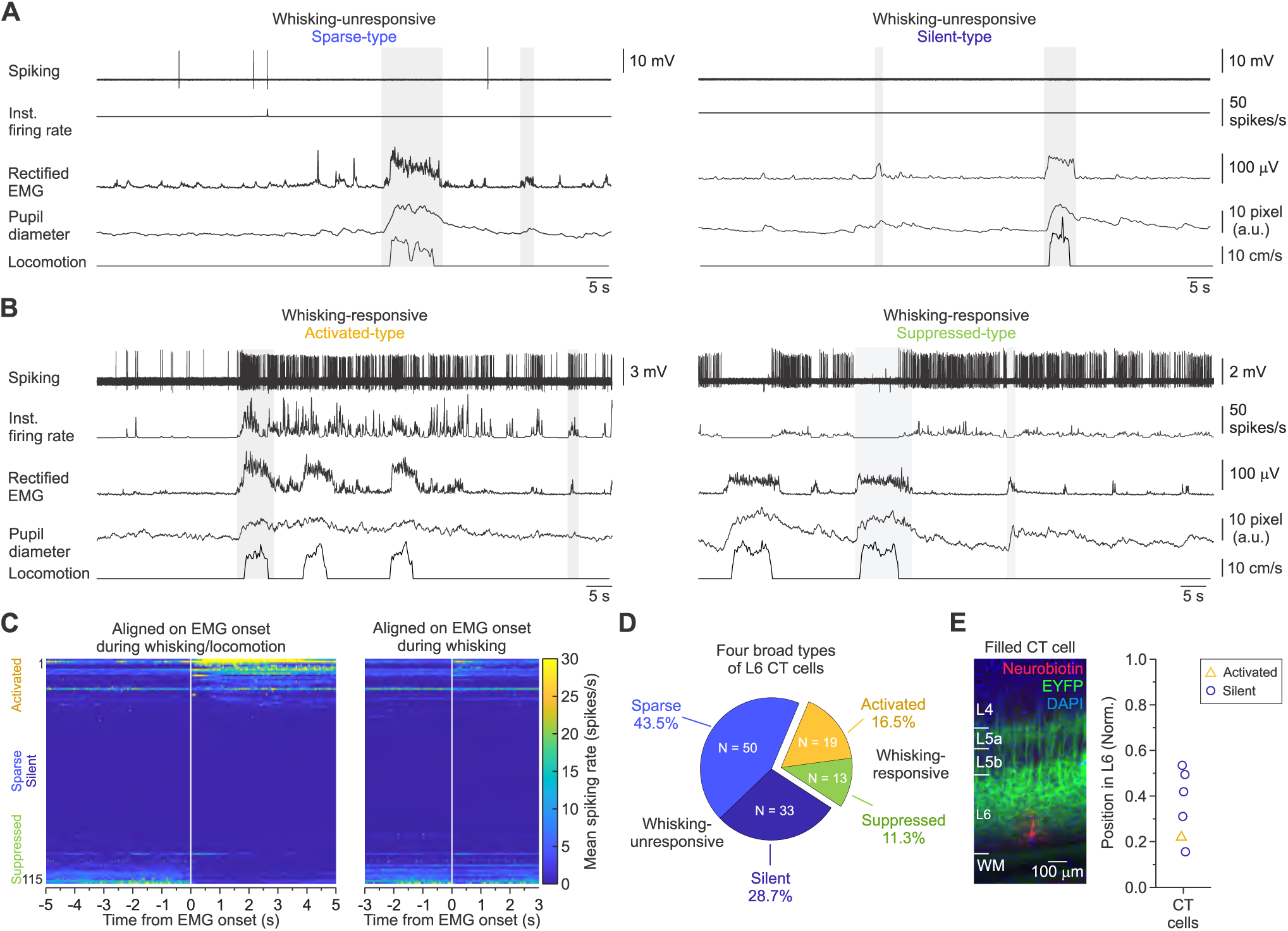
L6 CT neuron spiking as a function of behavioral state in awake, head-fixed mice. ***A,*** Representative data from two types of Whisking-unresponsive L6 CT neurons recorded during quiet awake periods and self-initiated whisking-related behaviors (two examples highlighted by grey shading): Sparse-type (left) and Silent-type (right). The mean firing rate for the Sparse-type CT cell was 0.012 spikes/s, whereas the Silent-type never fired during the entire recording session. Instantaneous firing rate is shown, together with rectified EMG of the contralateral mystacial pad musculature, pupil diameter, and walking patterns on the same time base. The instantaneous firing rate and pupil diameter were smoothed with a low pass Butterworth filter in both directions (100 ms window). ***B,*** Representative data from two types of Whisking-responsive L6 CT neurons recorded during quiet awake periods and self-initiated whisking-related behaviors (two examples highlighted by grey shading): Activated-type (left) and Suppressed-type (right). The Activated-type cell increased spiking rates during self-initiated whisking-related behavior, whereas the Suppressed-type cell decreased spiking rates. ***C,*** Overview of all recorded activity from L6 CT cells, sorted by firing rate difference between baseline (−4 to -1 s) and whisking (0 to 3 s) during both whisking/locomotion (left) and whisking only (right). The white line indicates EMG onset, and the color coding corresponds to the mean firing rate. ***D,*** Pie chart displaying the proportions of CT cell types identified based on their activity patterns. ***E,*** Left: Fluorescent image of a fixed slice (100 μm) showing a post hoc histologically verified ChR2-expressing CT cell within L6. Note the robust EYFP expression in L6 and a weaker fluorescence band near the L4-5 border. Right: Depth distribution of all histologically confirmed CT neurons.

The remaining L6 CT cells showed significant modulation in firing rates during either whisking or whisking/locomotion (Whisking-responsive: 28%; 32 of 115 CT neurons; mean spiking rate change around whisking onset; p < 0.05; Wilcoxon signed rank test; **Fig. 2*B*,*C,D***). Approximately 60% (19 of 32) of the Whisking-responsive CT cells had negligible firing rates during quiet wakefulness but were activated during whisking episodes, increasing their firing rates around solitary whisking, bouts of whisking/locomotion, or both (Activated-type). Conversely, roughly 40% (13 of 32) of the Whisking-responsive CT cells were spontaneously active during quiet wakefulness, but whisking and whisking/locomotion suppressed their spiking activity (Suppressed-type). Therefore, most L6 CT neurons in wS1 of awake mice are relatively unresponsive during whisking behavior, whereas approximately one-third of the cells showed distinct whisking-related modulations in firing rate (**Fig. 2*D***).

There was no clear relationship between relative depth in electrode penetration and the responsiveness of L6 CT cells. However, this could be due to an error in accurately localizing the pipette relative to the dural surface or the angle of the pipette not being perfectly normal to the cortical surface across penetrations. Nevertheless, 5 of the 6 CT neurons filled with neurobiotin were of the Silent-type, and their somata were located deeper within L6 (**Fig. 2*E***). The other was an activated CT neuron, and it was located within the lower third of the layer.

### Changes in L6 CT activity align better with whisking onset than locomotion but are more modulated during running

Consistent with previous observations (Sofroniew et al., 2014; Ayaz et al., 2019), mice never walked/ran on the wheel without whisking. To determine if the associated changes in L6 CT activity patterns are specific to whisker movement or locomotion, we examined the temporal dynamics of changes in firing around both whisking and locomotion onsets during bouts of walking/running (**Fig. 3*A,B*;** *see* Materials and Methods for the criteria used to define onset). These analyses revealed that Activated- and Suppressed-type CT cells’ firing rate changes aligned better temporally with EMG/whisking onset than locomotion (**Fig. 3*C***), suggesting locomotion does not activate most wS1 whisking-responsive L6 CT cells.

**Figure 3.**
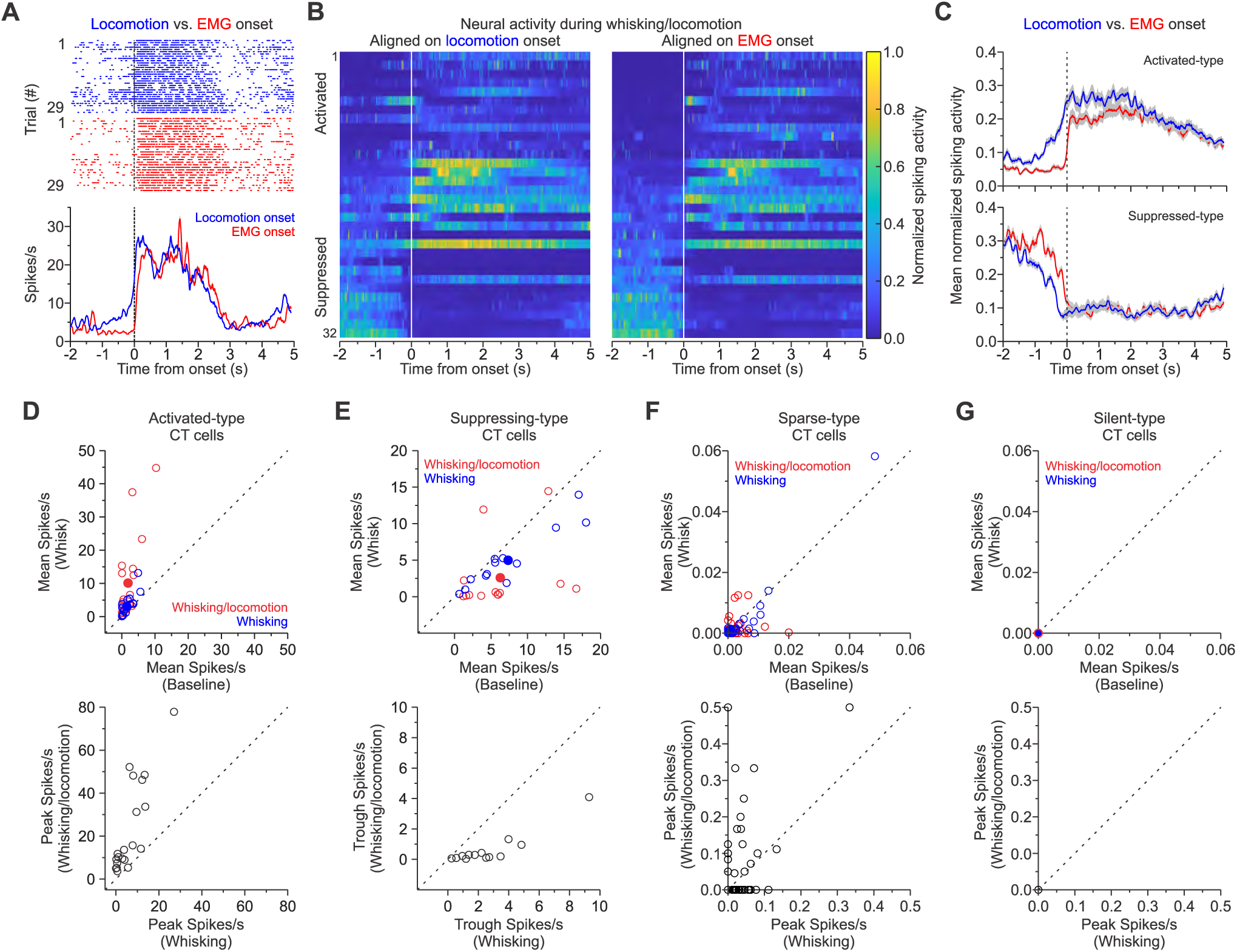
Changes in activity during whisking with concomitant locomotion align better with EMG onset than locomotion. ***A,*** Juxtacellular unit responses recorded from an example Activated-type CT neurons displayed as raster plots (top) and PSTHs (bottom) aligned on locomotion onset (blue raster and line) and EMG onset (red raster and; time = 0) during bouts of whisking/locomotion. Note the better temporal alignment between the change in firing rates and EMG onset than locomotion. ***B,*** Left: The normalized spiking patterns of all Whisking-responsive wS1 L6 CT neurons aligned on locomotion onset (left) or EMG onset (right) during whisking/locomotion. Each row is an individual neuron normalized to maximum firing rate and sorted by the mean baseline activity (3 s preceding EMG onset; white line; time = 0). ***C,*** Population PSTHs for both Activated- (n = 19 cells) and Suppressed-type CT cells (n = 13 cells) aligned on locomotion onset (blue line) and EMG onset (red line; time = 0). Neural activity is plotted as mean ± SEM (grey shadow indicating SEM). **D,** Summary graphs of spiking activity for all Activated-type CT cells. Top: Mean firing rate during whisking/locomotion (red circles) and solitary whisking (blue circles), plotted as a function of mean baseline activity preceding EMG onset (n = 19 cells; p = 2.44e-04 and 4.88e-04, respectively; Wilcoxon signed-rank test). Bottom: Peak firing rate for each neuron during whisking/locomotion plotted as a function of peak activity during whisking (p = 2.44e-04; Wilcoxon signed-rank test). **E,** Summary graphs of spiking activity for all Suppressed-type CT cells. Top: Mean firing rate during whisking/locomotion (red circles) and whisking (blue circles), plotted as a function of mean baseline activity preceding EMG onset (n = 13 cells; p = 0.03 and 4.88e-04, respectively; Wilcoxon signed-rank test). Bottom: Trough (lowest) firing rate for each neuron during whisking/locomotion plotted against that for whisking (p = 2.44e-04, Wilcoxon sign-rank test). **F-G,** Summary graphs of spiking activity for all Sparse- and Silent-type CT cells. Top: Mean firing rate during whisking/locomotion (red circles) and whisking (blue circles), plotted as a function of mean baseline activity preceding EMG onset (Sparse-type: n = 50, p = 0.1 and 0.09, respectively; Wilcoxon signed-rank test). Bottom: Peak firing rate for each neuron during whisking/locomotion plotted as a function of peak activity during whisking (Sparse-type: n = 50, p = 0.3; Wilcoxon signed rank test). The filled circles represent means in D-G. Values are represented as mean ± SEM.

Although the onset of locomotion does not necessarily activate most L6 CT neurons, the whisking-evoked response in these cells appeared to be more strongly modulated by locomotion. Therefore, we determined the cell-type-specific modulation of firing rates during whisking-related behaviors. On average, the mean firing rates of the Activated-type CT cells (n = 19) were low during baseline periods of quiet wakefulness before whisking-related behaviors (before whisking: 1.5 ± 1.7 spikes/s; before whisking/locomotion: 1.9 ± 2.6 spikes/s). During bouts of solitary whisking, the mean discharge rates of Activated-type cells were significantly higher than baseline (3.04 ± 3.23 spikes/s; p = 5.16e-04; Wilcoxon paired signed-rank test; **Fig. 3*D***, top blue circles). Responses were also significantly elevated during whisking/locomotion, becoming more intense, with some having high firing rates (10.12 ±12.2 spikes/s; p = 8.85e-05; Wilcoxon paired signed-rank test; **. 3*D***, top red circles). The peak firing rates across all Activated-type CT cells sampled were significantly higher during whisking/locomotion (22.87 ± 21.16 spikes/s) than whisking (6.5 ± 6.7 spikes/s) (p = 1.033e-04; Wilcoxon paired signed-rank test, **Fig. 3*D***, bottom).

All Suppressed-type CT cells (n = 13) were spontaneously active during quiet wakefulness preceding whisking-related behaviors (before whisking: 6.3 ± 5.2 spikes/s; before whisking/locomotion: 7.3 ± 5.6 spikes/s). However, the mean firing rate of these neurons significantly decreased during whisking (5.0 ± 4.0 spikes/s; p = 1.03e-04; Wilcoxon paired signed-rank test; **Fig. 3*E***, top blue circles) and whisking/locomotion (2.6 ± 4.8 spikes/s; p = 0.03; Wilcoxon paired signed-rank test; **Fig. 3*E***, top red circles). The three Suppressed-type cells with a mean firing rate higher during whisking/locomotion (red data points above the diagonal) correspond to those that responded bimodally, with an initial decrease followed by an increase in spiking activity during locomotion epochs. Across all Suppressed-type CT cells recorded, the lowest level (trough) of instantaneous spike rate during whisking/locomotion (0.6 ± 1.1 spikes/s) was lower than that observed for whisking (2.7 ± 2.4 spikes/s; p = 2.44e-04; Wilcoxon paired signed-rank test; **Fig. 3*E***, bottom). As expected, the mean firing rates of both Sparse- and Silent-type CT cells (n = 83) were low during baseline periods of quiet wakefulness before whisking-related behaviors (most < 0.02 spikes/s for Sparse-type) and did not change significantly during solitary whisking or whisking/locomotion (p > 0.05; Wilcoxon paired signed-rank test; **Fig. 3*G***).

In summary, Activated- and Suppressed-type L6 CT cells recorded in wS1 exhibited robust modulation of firing rates during whisking-related behavior, with changes in firing rate better aligned with EMG than locomotion. However, these same neurons exhibited more significant firing rate modulation during whisking/locomotion than during bouts of whisking only.

### Diverse response types of L6 CT cells during whisker-related behaviors

Further examination of the whisking-responsive CT population revealed key relationships between behavior and neuronal firing rate. Specifically, we found that Activated-type cells showed diverse responses and significant modulation of spiking rates during whisking/locomotion that comprised tonic increases in spike rate, phasic increases in the rate at whisking onset, elevated firing rates before whisking onset, and delayed activity (**Fig. 4*A***, Activated-type, from left to right). In addition, most Activated-type cells (68%; 13 of 19) increased spiking during solitary whisking, with the only exception being those that had delayed activity during whisking/locomotion (6 of 19 cells; **Fig. 4*B***). These cells suggest changes in firing rates of some Activated-type neurons are not always associated with whisking onset. However, as illustrated in **Figure 4A** (the fourth cell to the right), the changes in spiking did not align with locomotion onset either. Suppressed-type CT cells exhibited less diverse responses, with most responding with a decrease in spike rate during whisking/locomotion and whisking (**Fig. 4*A,B***). However, we did encounter three Suppressed-type cells with more complex firing patterns, such as responding bimodally, with first a decrease and then an increase in firing rate during locomotion.

**Figure 4.**
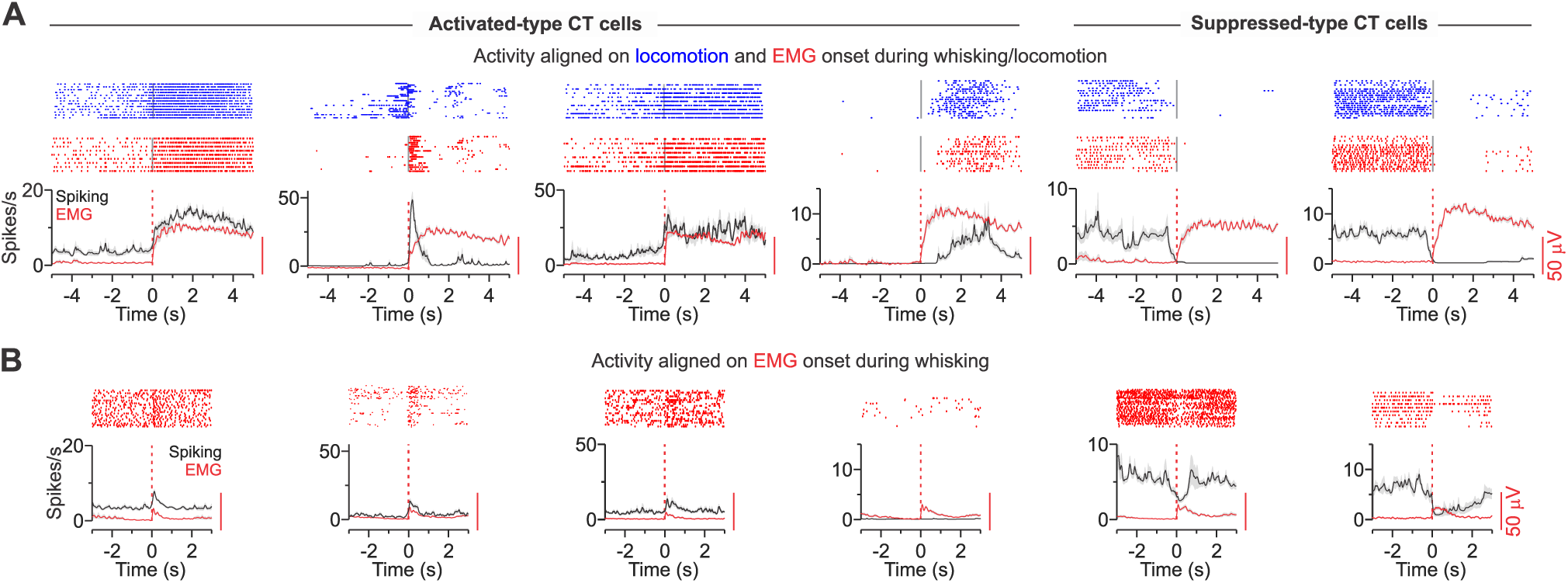
Diverse response profiles exhibited by whisker-related L6 CT neurons in wS1. ***A,*** *Top:* Examples of juxtacellular unit responses recorded from individual L6 CT neurons displayed as raster plots aligned on locomotion onset (upper row; blue) and EMG onset (lower row; red dashed line; time = 0) during whisking/locomotion. *Below:* PSTHs from the same cell aligned to EMG onset. The first four examples (left to right) show Activated-type CT neurons, whereas the last two are Suppressed-type CT cells. Activated-type cells had either tonic firing at EMG onset (left), phasic increases in activity at EMG onset (second from left), a slow build-up of activity preceding EMG onset (third from left), or delayed increases in activity from EMG onset (fourth from left). Suppressed-type neurons showed less diversity and tended to have decreased spiking rates before EMG onset. For the PSTHs, neural activity (black line) and EMG activity (red line) are plotted as mean ± SEM (grey shadow indicating SEM). ***B,*** Juxtacellular unit responses for the same CT neurons (shown in A) displayed as raster plots and PSTHs aligned on EMG onset (red dashed line; time = 0) during whisking and no locomotion. We found that most CT cells were modulated during solitary whisking similarly to whisking/locomotion. The only exception were cells that had delayed increases in activity during locomotion (i.e., the fourth cell from left), which did not respond during whisking only. Values are represented as mean ± SEM.

### Pre-movement modulation of firing rate in Whisking-responsive L6 CT cells of wS1

For many Whisking-responsive L6 CT neurons, we often observed changes in spiking rates that preceded whisking behavior, as measured by the electrical activity generated in the muscles of the mystacial pad (i.e., EMG signal). To assess the temporal relationship between whisker movement and a change in neural activity, we characterized the firing rate changes in Whisking-responsive L6 CT neurons (Activated- and Suppressed-type) as a function of time around the onset of self-initiated whisking. Since we were interested in changes in spiking activity around EMG onset, we excluded the Activated-type cells with delayed activity during locomotion from subsequent analyses (n = 6 of 19 Activated-type cells). We defined a change in instantaneous spiking activity as a 30% deviation (above or below) from the average pre-whisking baseline activity (2 s). This criterion was conservative and biased towards delaying onset detection. Since the analysis depended on baseline activity, we selected only neurons with stable rates above 2.0 spikes/s. Among all the Activated- and Suppressed-type CT cells recorded, 38% (5 of 13) and 77% (10 of 13) met this criterion for analysis, respectively.

**Figure 5*A*** shows two Activated- and Suppressed-type cells with a firing rate change (black dotted line) detected before EMG onset (red dotted line). Of the five Activated-type cells analyzed, 80% (4 of 5 cells) exhibited rate changes that increased before whisker movement (mean: -118 ± 178 ms, n = 5 cells, **Fig. 5*B***, top). Similarly, the discharge rate in 90% (9 of 10 cells) of Suppressed-type cells decreased before EMG onset (mean: -248 ± 206 ms, n = 10 cells, **Fig. 5*B***, bottom). Since we performed the above analysis on whisking/locomotion, we repeated the analysis on bouts of solitary whisking (**Fig. 5C**). Due to the variability in EMG signal related to whisking, we focused our analysis on the strongest 50% of all whisks identified for each cell based on the peak EMG signal. Most Activated-and Suppressed-type cells showed similar spiking rate changes before solitary whisking. Altogether, these data suggest that a subset of L6 CT cells in wS1 change their firing rate well before self-initiated whisker movements, with and without subsequent locomotion.

**Figure 5:**
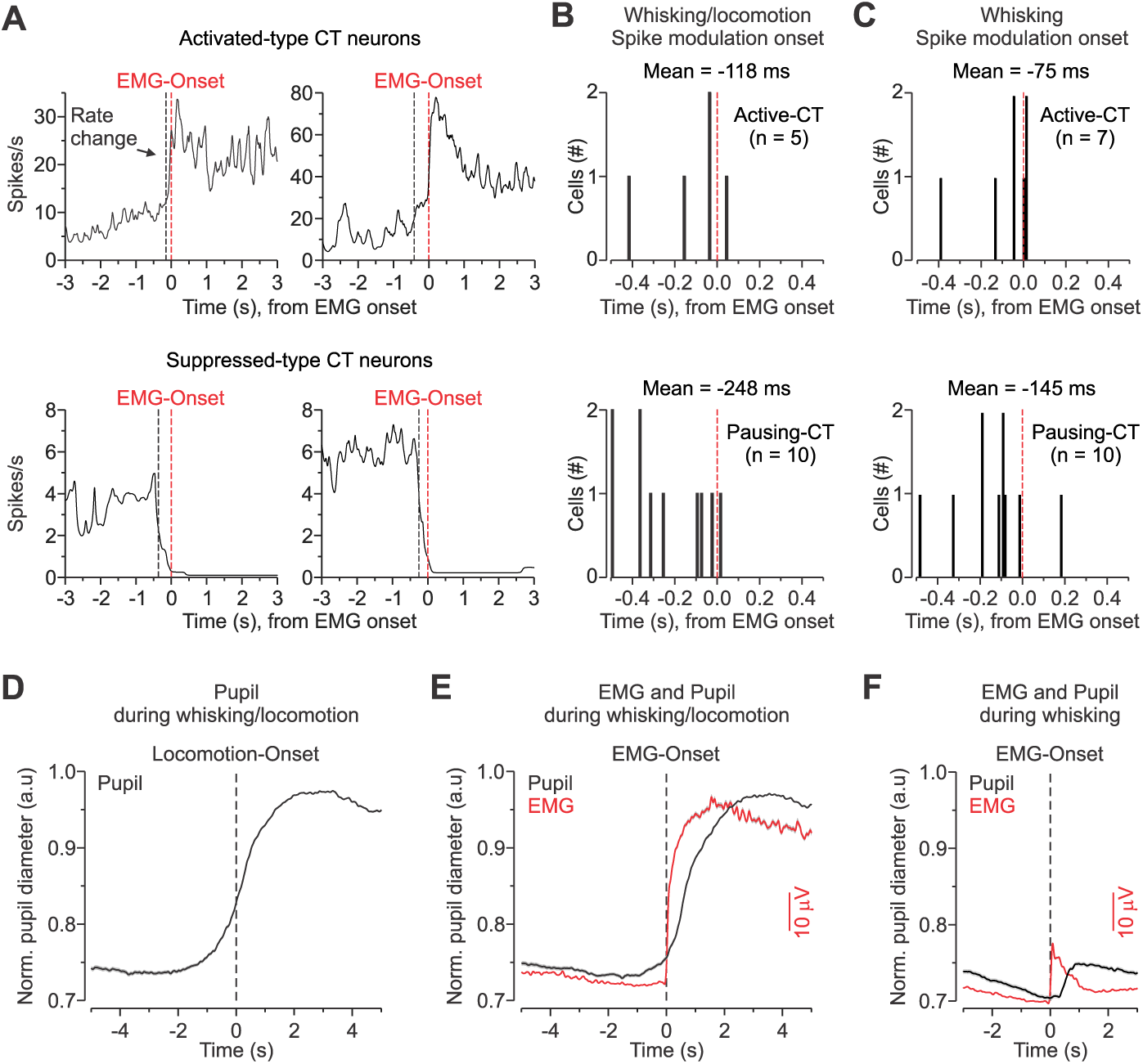
Subpopulations of Whisking-responsive L6 CT cells are modulated before whisker movement. ***A,*** Top: Two Activated-type CT neurons exhibiting pre-EMG spike enhancement. Bottom: Two Suppressed-type CT neurons exhibiting pre-EMG spike suppression. The red and black vertical dashed lines indicate EMG onset and the detected whisker-related spike modulation, respectively. ***B,*** Shown are the onset latencies of the pre-EMG spiking modulation for 5 Activated- and 10 Suppressed-type CT neurons with a mean baseline firing rate greater than 2 spikes/s before whisking/locomotion. Most Activated-type CT cells (8 of 13) had little or no baseline firing rate, rendering them unsuitable for analysis (see text), whereas most Suppressed-type CT neurons (9 of 10) exhibited pre-EMG spiking suppression. ***C,*** Shown are the onset latencies of the pre-EMG spiking modulation for 7 Activated- and 10 Suppressed-type CT neurons with a mean baseline firing rate greater than 2 spikes/s before solitary whisking episodes. Like B, most Suppressed- and some Activated-type CT neurons exhibited pre-EMG spiking modulation. ***D,*** Averages of mean normalized pupil diameter and mean locomotion activity of all locomotion bouts for each neuron. Data is aligned on the onset of locomotion across all sessions when a Whisking-responsive neuron was recorded (n = 32 sessions). Pupil diameter was normalized to maximum dilation during the session. Values are represented as mean ± SEM. ***E,*** Averages of mean EMG and mean pupil diameter during whisking/locomotion (same data shown in D) but now aligned on EMG onset. Values are represented as mean ± SEM. ***F,*** Averages of mean EMG and mean normalized pupil diameter during whisking aligned on EMG onset across all sessions (n = 32 sessions). Pupil modulation occurs after EMG onset. Values are represented as mean ± SEM.

The transition from quiet awake to exploratory behaviors such as whisking and locomotion is associated with changes in arousal (McGinley et al., 2015b). To help understand the contributions of arousal to the observed changes in L6 CT firing rates, we analyzed changes in pupil diameter, which is a reliable index for general arousal in head-fixed mice (Reimer et al., 2014; McGinley et al., 2015a; Vinck et al., 2015). Consistent with these previous studies, we observed that the onset of locomotion is preceded by an increase in pupil diameter, which ramped up hundreds of milliseconds before, suggesting that general arousal may contribute to the pre-movement modulation of CT activity (n = 32 behavioral sessions obtained from the 32 whisker-related CT cells; **Fig. 5*D***). However, while pupil diameter increased 500-1000 ms before EMG onset during whisking/locomotion, we found that during solitary whisking, the increases in the EMG signal preceded the change in pupil diameter by 250-500 ms (**Fig. 5*E-F***). These results suggest that the pre-movement modulation of L6 CT circuit activity may also represent a motor-preparatory signal.

To assess the contributions of arousal and motor activity to the cell-type-specific modulation of spiking rate observed in Whisking-responsive L6 CT cells, we calculated the cross-correlation coefficient between EMG signal or pupil diameter and the instantaneous spiking rate around whisking onset. Analysis of the time courses of EMG signals and pupil dynamics revealed a moderately strong correlation and anti-correlation with spiking activity for most Activated- and Suppressed-type CT cells, respectively, during both whisking/locomotion and whisking (**Fig. 6*A,B***). Overall, we found that the best absolute cross-correlation (irrespective of lags or leads) for both Activated- and Suppressed-type cells was consistently stronger with EMG than with pupil during whisking/locomotion and whisking (Activated-type CT cells: during whisking/locomotion: p = 0.06; whisking: p = 2.44e-04; Suppressed-type CT cells: during whisking/locomotion: p = 0.002; whisking: p = 0.27; Wilcoxon paired signed-rank test; **Fig. 6*C,D***, left). Moreover, we found that EMG signals generally aligned with the spiking activity of Whisking-responsive CT cells with a much shorter lag than pupil dynamics (Activated-type CT cells: during whisking/locomotion: p = 0.002; whisking: p = 2.44e-04; Suppressed-type CT cells: during whisking/locomotion: p = 0.002; whisking: p = 0.08; Wilcoxon paired signed-rank test; **Fig. 6*C,D***, right). Altogether, these results indicate a stronger correlation and better temporal alignment between the firing rates of Whisking-responsive CT cells and the time course of the EMG (motor) signal than pupil (arousal) dynamics.

**Figure 6.**
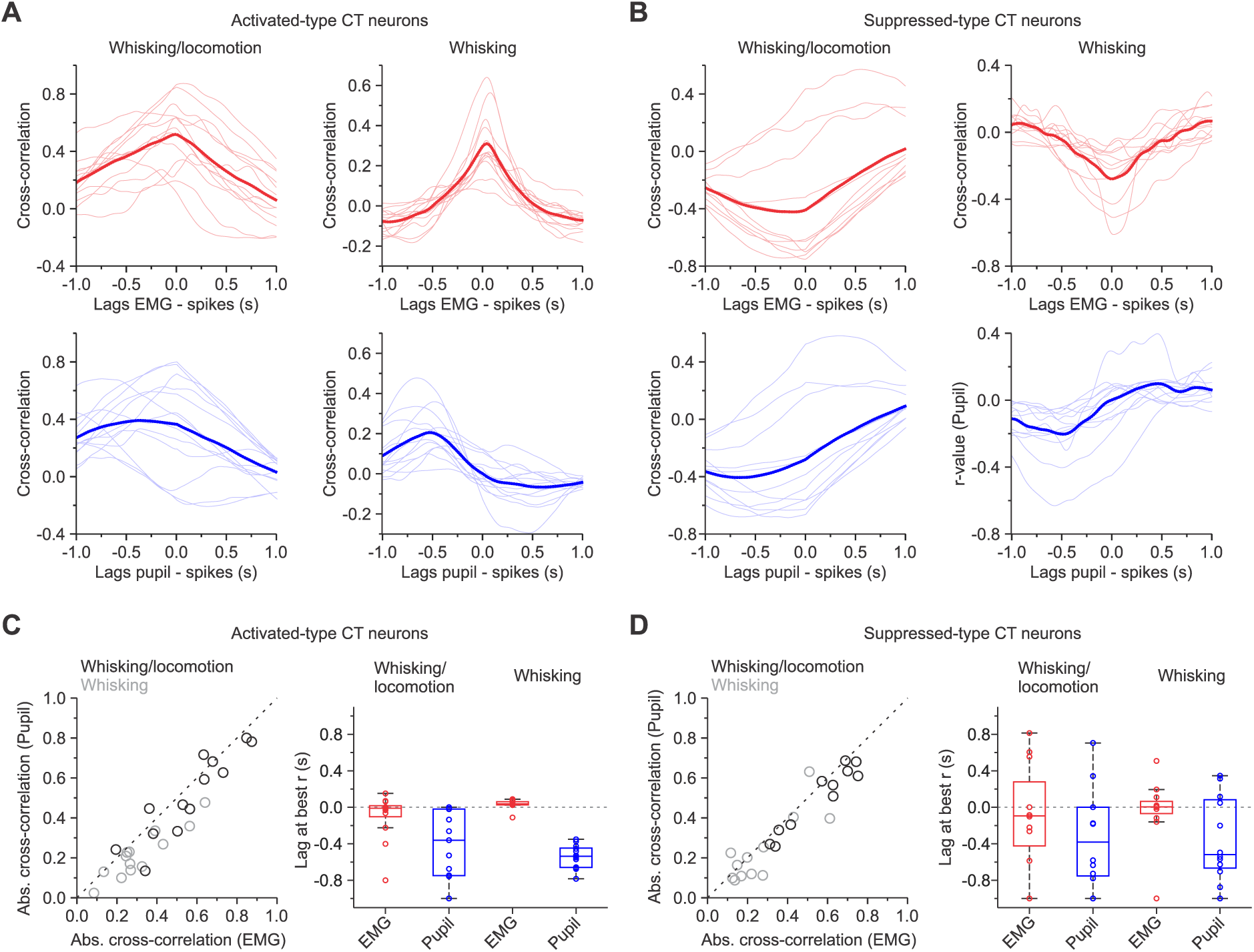
Contributions of whisking and arousal to L6 CT neuron in wS1 of awake mice. ***A-B,*** Cross-correlograms of spiking activity and EMG signals (top, red) or pupil dynamics (bottom, blue) from Activated-type (***A***) and Suppressed-type CT neurons (***B***) during whisking/locomotion (left) and whisking (right). Light-colored lines (red and blue) represent the average trial-by-trial cross-correlations for each Whisking-responsive CT neuron. The dark-colored lines (red and blue) represent the mean cross-correlation across all neurons (n = 13/13 Activated-type cells; n = 11/12 Suppressed-type cells for whisking/locomotion and whisking, respectively). ***C-D,*** Left, summary graphs showing the absolute peak cross-correlation of pupil dynamics to firing activity as a function of the peak cross-correlation of EMG and firing activity for each Activated- (***C***, left) and Suppressed-type CT cell (***D***, left) during whisking/locomotion (black circles) and whisking (grey circles). Right, summary graphs showing the comparison of lags at which peak absolute correlation coefficients were obtained for Activated- (***C,*** right) and Suppressed-type CT neurons (***D,*** right) during whisking/locomotion and whisking (Activated-type mean cross-correlation values: EMG during whisking/locomotion = 0.56 ± 0.20; Pupil during whisking/locomotion = 0.51 ± 0.21; EMG during whisking = 0.31 ± 0.16; Pupil during whisking = 0.22 ± 0.12) (Suppressed-type mean cross-correlation values: EMG during whisking/locomotion = 0.56 ± 0.17; Pupil during whisking/locomotion = 0.50 ± 0.16; EMG during whisk-only = 0.27 ± 0.16; Pupil during whisk-only = 0.23 ± 0.17). Data in ***C*** (right) and ***D*** (right) are plotted as median, interquartile distance, and range. Individual data points are represented as open circles.

### Relationship between pupil diameter and spiking activity of Whisking-unresponsive CT cells

Like S1 CT cells in other species (Landry and Dykes, 1985; Swadlow, 1989, 1990; Swadlow and Hicks, 1996; Kelly et al., 2001; Lee et al., 2008; Kwegyir-Afful and Simons, 2009), most L6 CT cells recorded in the present study showed very little spontaneous activity or whisker-related responses (72%; **Fig. 2*A***). Here, we routinely observed that Whisking-unresponsive CT cells with low spontaneous firing (i.e., Sparse-type) exhibited a strong relationship to the moment-to-moment fluctuations in pupil diameter. **Figure 7*A*** shows an example of a neuron with a spontaneous rate of 0.6 spikes/s. To analyze this relationship, we calculated the spike-triggered average of the pupil diameter and found that this cell routinely spiked during constriction events (362 total spikes during a ∼10 min recording period; **Fig. 7*B***). Across 43 Sparse-type CT cells that discharged a minimum of 5 spikes during the recording session, nearly all (91%; 39 of 43) showed a similar relationship between individual spikes and pupillary constriction, as verified by the negative pupil slope (**Fig. 7*C***). The average spike-triggered changes in pupil diameter from all recorded units indicate a similar effect, with spikes occurring during constriction events (n = 5243 spikes from 50 Sparse-type CT cells, recorded over 29090 s; **Fig. 7*D***). These data indicate that most spiking activity in Sparse-type CT cells occurs during pupillary constriction. Interestingly, the relationship between the change in pupil diameter and Sparse-type spiking was independent of the waking state, as we found qualitatively similar patterns of effects whether mice were sitting still (quiet) or actively exhibiting exploratory movements (**Fig. 7*E***; i.e., whisking or whisking/locomotion).

**Figure 7.**
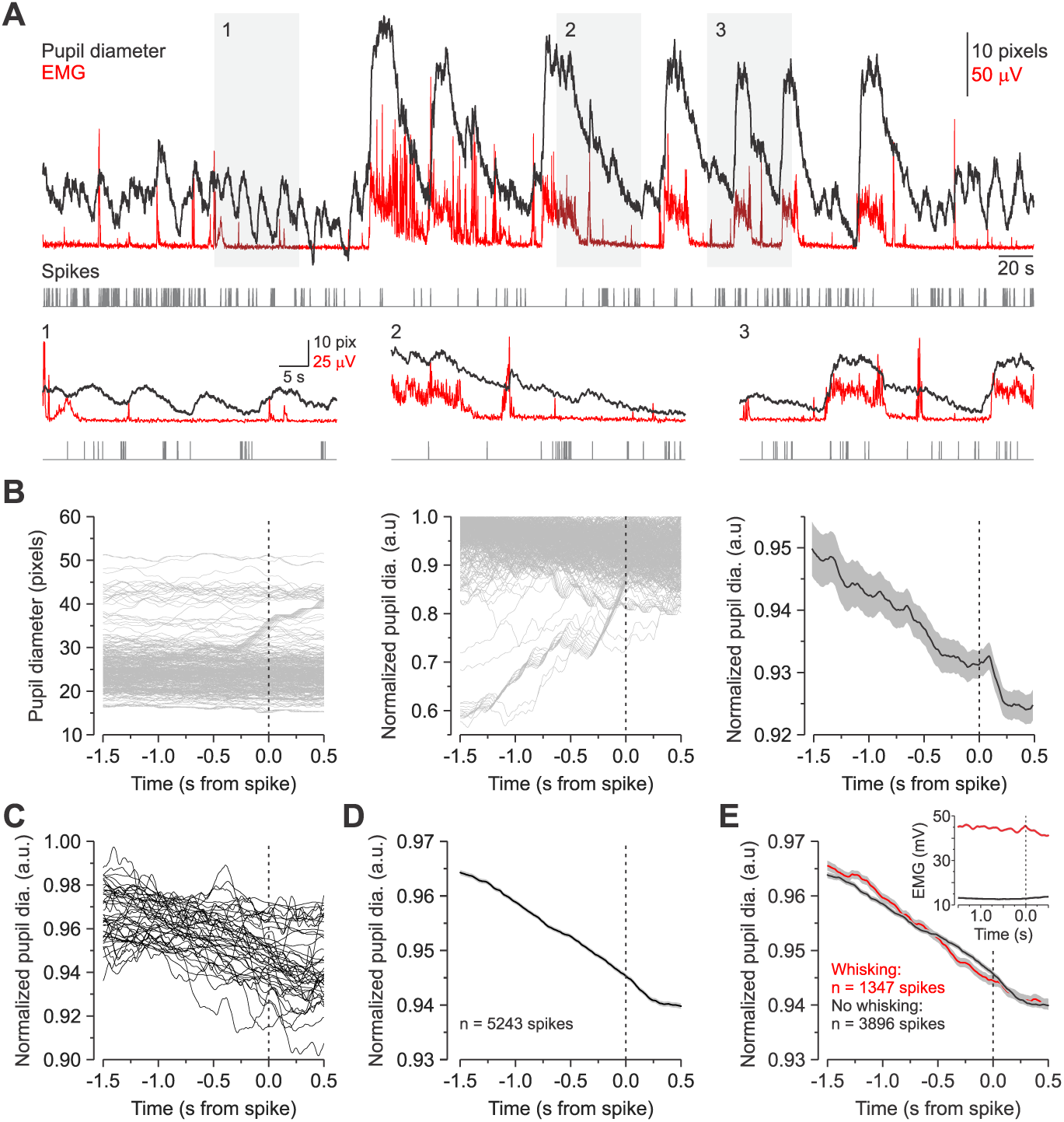
Whisking-unresponsive L6 CT cells with sparse firing preferentially fired during pupil constrictions. ***A,*** Pupil diameter (black trace), rectified EMG (red trace), and spike times (grey ticks) are shown as a function of recording time for a Sparse-type CT cell. Insets, expansion in time showing the relationship between spiking, EMG, and pupil diameter (1, 2, and 3; grey shading). Note small increases in pupil diameter are seen even in the absence of whisker movement (i.e., EMG changes). Note the increased incidence of spikes during pupil constriction. ***B,*** Spike-triggered pupil average for the same pupil cell pair (shown in **A**). Left: Raw pupil traces aligned on spike onset (n = 362 spikes; time = 0, dotted vertical line). It is hard to appreciate any trends in pupillary responses as they spanned a broad range of pupil diameters. For better visualization, we normalized the extracted pupillary responses to the peak of each extracted pupillary response. Middle: The same spike-aligned pupil traces normalized to their peak value. Note the clear tendency of preceding pupil constriction in most instances just before spiking. Right: Mean and SEM of the normalized spike-triggered pupil (black line and grey shadow). ***C,*** The normalized spike-triggered pupil average for each Sparse-type CT neuron (n = 43/50 cells; similar to B, right panel). Only Sparse-type cells with more than five spikes recorded in the entire session were used for analysis (7 cells excluded with < 5 spikes). ***D,*** A summary graph of all normalized spike-triggered pupil traces (n = 5243 traces) across all the silent CT cells (n = 50 cells). ***E,*** A summary graph showing the spike-triggered pupil traces shown in D after separating them based on EMG activity at the time of the spike. Inset, the mean spike-triggered EMG signal across the two behavioral states (Whisking/high EMG activity, n = 1347 spikes; No whisking/low EMG activity, n = 3896 spikes). Whisking/high EMG activity was defined as an average EMG exceeding 20 μV in the 1.5 s preceding the spike. Values are represented as mean ± SEM.

## DISCUSSION

In this study, we have used the juxtacellular method to record optogenetically identified L6 CT neurons in wS1 of awake, head-restrained mice during quiescence and whisking-related behavior. We characterized the modulation of CT spiking across different behavioral states through continuous monitoring of arousal, whisking, and locomotion, making several novel observations. First, we found two subsets of CT cells that exhibited opposite spiking rate changes as mice transitioned from quiet wakefulness to whisking: Activated and Suppressed. Second, some CT neurons, primarily the Suppressed-type, presented changes in firing rate well before whisking onset. Third, while arousal and whisking contribute to state-dependent changes in firing rates, the activity of Whisking-responsive cells correlated more strongly and displayed better temporal alignment with whisking onset. Fourth, most L6 CT neurons in wS1 are spontaneously quiet (Sparse-type) or entirely silent in the awake mouse, even during active whisking and locomotion. Finally, if the Sparse-type cells spike, they preferentially fired at the state transition point corresponding to when pupil diameter slowly constricted and the mouse entered a period of quiet wakefulness. Thus, our results demonstrate that L6 CT neurons in wS1 of behaving mice have different response properties and precisely timed spikes around distinct state transition points.

### Whisking-responsive L6 CT neurons

28% of L6 CT neurons encountered exhibited responses to whisking-related behavior. Because of their response properties, we differentiated this population into two types, an Activated-type with low spontaneous rates that increased spiking around whisking and a Suppressed-type with moderate spontaneous rates that decreased spiking around whisking. These response types are qualitatively similar to the 25% of wS1 CT cells reported to be either excited or inhibited by controlled whisker deflection in the narcotized rat (Kwegyir-Afful and Simons, 2009). The complimentary activity patterns are also reminiscent of recent findings in other sensory cortices of *Ntsr1-Cre* mice during awake behavior (Augustinaite and Kuhn, 2020; Clayton et al., 2021), suggesting a general feature of CT processing. However, naturally occurring transitions between distinct motor behaviors - whisking and locomotion - revealed heterogeneity within wS1. For example, we observed both phasic and tonic changes in firing rate during whisking associated with locomotion. Since mice begin whisking shortly before locomotion onset, phasic responses suggest that whisking and locomotion signals may have opposing effects on some CT cells. We also encountered a small subset of neurons active only during locomotion, yet locomotion does not activate these cells per se since their change in spiking activity was not well aligned with the onset of locomotion. Together, these results confirm that locomotion contributes to response diversity in wS1 (Sofroniew et al., 2015; Ayaz et al., 2019). Although they represent a minority of CT cells, these distinct Whisking-responsive populations appear well-suited to dynamically regulate thalamocortical processing during sensorimotor behavior (Steriade, 2001).

One function of these CT cells may be to selectively enhance or suppress afferent sensory signals in a context-dependent manner. CT fibers from L6 are glutamatergic and thus a direct source of monosynaptic excitatory input to thalamic relay cells (Scharfman et al., 1990; Bourassa et al., 1995; Golshani et al., 2001), but they also indirectly inhibit them by recruiting GABAergic neurons in the thalamic reticular nucleus (TRN) (Cox et al., 1997; Kim et al., 1997; Golshani et al., 2001; Crandall et al., 2015). Previous work has shown that the sign of CT influence on thalamic relay cells is highly dynamic, with the excitatory-inhibitory balance shifting in an activity-dependent manner (von Krosigk et al., 1999; Crandall et al., 2015). CT effects are primarily suppressive during low-frequency activity, whereas higher-frequency activity yields enhancement. Consistent with the findings of Augustinaite and Kuhn (2020), we find that the activity of Activated- and Suppressed CT cells complement each other according to behavioral state, suggesting they may also have complementary effects on thalamic excitability. For example, when Activated-type CT cells are firing at high rates during whisking, Suppressed-type cells are inhibited, and when Activated-type cells are firing at low frequencies in the absence of whisking, Suppressed-type cells spike at higher rates. Given that L6 CT cells have diverse termination patterns across and within thalamic nuclei (Deschenes et al., 1998), future studies identifying the target of these CT cells will help understand how these circuits control the spatial and temporal structure of thalamocortical activity during awake behavior (Temereanca and Simons, 2004; Crandall et al., 2015; Whitmire et al., 2016; Martinez-Garcia et al., 2020).

### Premovement modulation of L6 CT activity

This study found that some Activated- and most Suppressed-type CT cells showed significant response modulation well before whisking onset. What might be the source of this premovement modulation in CT activity? One possibility is that the change in activity may involve recruiting neuromodulatory systems that control global arousal levels (Lee and Dan, 2012). Indeed, exploratory behaviors such as whisking and locomotion are closely associated with arousal (McGinley et al., 2015b). Here we monitored pupil diameter to assay arousal (Reimer et al., 2014; McGinley et al., 2015a; Vinck et al., 2015), finding a strong correlation between the firing rates of Whisking-responsive CT neurons and the time course of pupil dynamics. Consistent with visually evoked responses in L6 CT cells of the visual cortex (Augustinaite and Kuhn, 2020), we found that the average spiking activity of Whisking-responsive cells is more modulated during whisking/locomotion, a substate of heightened arousal, than whisking alone. However, while an increase in pupil diameter preceded whisking/locomotion, the increase in pupil diameter occurred after the onset of a solitary whisk, suggesting that the premovement modulation of L6 CT activity may also represent a motor-preparatory signal. Consistent with this interpretation, we found stronger correlations and better temporal alignment between the firing rates of Whisking-responsive cells and the time course of the EMG signal than pupil dynamics. Recently Clayton et al. (2020) reported that L6 CT activity in the auditory cortex is also modulated hundreds of milliseconds before orofacial facial movements associated with sound presentation and reward, but not locomotion. Together, these studies suggest that encoding preparatory activity is perhaps a general principle underlying the activity of some CT cells in sensory cortices. Such precisely timed changes in CT cell activity before self-generated whisker movement could provide context-dependent control of information processing in somatosensory thalamocortical circuits during active sensation (Lee et al., 2008). Indeed, previous studies in rodents have shown that L6 wS1 neurons receive excitatory input from the whisker primary motor cortex (Zhang and Deschenes, 1998; Lee et al., 2008; Rocco-Donovan et al., 2011; Kinnischtzke et al., 2014; Kinnischtzke et al., 2016; Martinetti et al., 2022), a cortical area that appears to contribute to the initiation and modulation of whisking (Carvell et al., 1996; Sreenivasan et al., 2016). The enhancing and suppressive effects of such motor-preparatory signals on CT firing most likely depend on the engagement of specific excitatory and inhibitory mechanisms in L6 (Beierlein and Connors, 2002; Crandall et al., 2017; Frandolig et al., 2019; Martinetti et al., 2022).

### Whisking-unresponsive L6 CT neurons

Most L6 CT neurons (72%) in mouse wS1 exhibited no whisking/locomotion-related activity and had either a low rate of spontaneous activity (Sparse-type) or were completely silent (Silent-type). This proportion of unresponsive cells is consistent with previous studies of antidromically identified CT neurons in S1 of sedated or anesthetized rats and awake rabbits (Landry and Dykes, 1985; Swadlow, 1989, 1990; Swadlow and Hicks, 1996; Kelly et al., 2001; Lee et al., 2008; Kwegyir-Afful and Simons, 2009), but is higher than those reported for other cortices (Tsumoto and Suda, 1980; Swadlow, 1994; Beloozerova et al., 2003; Sirota et al., 2005; Stoelzel et al., 2017; Augustinaite and Kuhn, 2020). Overall, there is considerable evidence across species and sensory modalities that many L6 CT neurons are fairly silent and unresponsive, even when animals are aroused and behaving, suggesting they are a common feature of CT systems. Yet, the function of this largely quiet population is a mystery.

We found Whisking-unresponsive cells with low spontaneous activity, or Sparse-type CT neurons, preferentially fired when the pupil diameter slowly constricted. Previous studies have shown that pupil constriction is associated with a decrease in cortical activation and an increase in the prevalence of low-frequency cortical oscillations (Reimer et al., 2014; McGinley et al., 2015a; Vinck et al., 2015), suggesting that some CT neurons themselves may play a role in switching cortical network activity. So how does the sparse but precisely timed activity of most excitatory L6 CT neurons impact cortical network activity? One possibility is synchronizing the output of translaminar L6 inhibitory cells (Olsen et al., 2012; Bortone et al., 2014), which could fire synchronously enough to reduce synaptic activity across all cortical layers, terminating cortical activation (Timofeev et al., 2001; Volgushev et al., 2006). Alternatively, synchronized low-frequency CT spiking could also suppress the thalamus through the preferential recruitment of inhibitory TRN neurons (Crandall et al., 2015).

Of course, the small behavioral space in which mice were allowed to rest quietly or self-initiate whisking and locomotion may have contributed to the large fraction of Whisking-unresponsive CT cells encountered in the present study. Indeed, evidence suggests that many unresponsive CT neurons have subthreshold excitatory and inhibitory receptive field properties (Swadlow and Hicks, 1996; Kwegyir-Afful and Simons, 2009). Thus, it seems reasonable that different behaviors, tasks, sensory stimuli, or brain states could bring these unresponsive CT cells online. Consistent with this idea, a recent imaging study by Augustininaite & Kuhn (2020) demonstrated that quiet L6 CT neurons in the visual cortex are recruited dynamically under different stimulation conditions or behavior states. Further investigations of the activity of wS1 L6 CT cells in mice performing different whisker-dependent tasks or transitioning between different brain states (waking versus sleep) could reveal additional insights into the functional significance of this mysterious population.

Indeed, considerable literature examining CT circuits describes diverse roles for L6 neurons in modulating sensory transmission. These functions include regulation of gain or excitability (Olsen et al., 2012; Denman and Contreras, 2015; Guo et al., 2017; Wang et al., 2018; Ansorge et al., 2020), sharpening or shifting of sensory response properties (Temereanca and Simons, 2004; Andolina et al., 2007; Li and Ebner, 2007; Hasse and Briggs, 2017; Pauzin and Krieger, 2018; Pauzin et al., 2019; Voigts et al., 2020), enhancement of synchrony (Sillito et al., 1994; Contreras et al., 1996), regulation of sensory adaptation (Mease et al., 2014), and control of burst vs. tonic spiking (Wang et al., 2006; Kirchgessner et al., 2020). The current work complements these studies, and, together, they indicate that L6 CT cells are highly heterogeneous and likely have multiple modulatory functions.

